# Design of a symmetry-broken tetrahedral protein cage by a method of internal steric occlusion

**DOI:** 10.1101/2023.11.08.566319

**Authors:** Nika Gladkov, Elena A. Scott, Kyle Meador, Eric J. Lee, Arthur D. Laganowsky, Todd O. Yeates, Roger Castells-Graells

## Abstract

Methods in protein design have made it possible to create large and complex, self-assembling protein cages with diverse applications. These have largely been based on highly symmetric forms exemplified by the Platonic solids. Prospective applications of protein cages would be expanded by strategies for breaking the designed symmetry, *e*.*g*., so that only one or a few (instead of many) copies of an exterior domain or motif might be displayed on their surfaces. Here we demonstrate a straightforward design approach for creating symmetry-broken protein cages able to display singular copies of outward-facing domains. We modify the subunit of an otherwise symmetric protein cage through fusion to a small inward-facing domain, only one copy of which can be accommodated in the cage interior. Using biochemical methods and native mass spectrometry, we show that co-expression of the original subunit and the modified subunit, which is further fused to an outward-facing anti-GFP DARPin domain, leads to self-assembly of a protein cage presenting just one copy of the DARPin protein on its exterior. This strategy of designed occlusion provides a facile route for creating new types of protein cages with unique properties.

## INTRODUCTION

Advances in protein design have made it possible, with sufficient experimental trials, to generate self-assembling protein nanoparticles or protein cages, which often take the form of Platonic solids (tetrahedra, cubes, icosahedra) ^1–17^. Principles of symmetry have played a critical role in successful strategies for designing protein cages and similar assemblies ^16,18–20^. Building symmetric structures minimizes the engineering requirements for achieving robust assembly. That principle was first emphasized by Crick and Watson ^21^ in the context of natural assemblies (viral capsids) and was a guiding principle in developing the first method for designing novel protein cage assemblies ^14^.

A consequence of design methods that exploit symmetry is that the resulting architectures are repetitive. For example, the outer surface of a symmetric protein cage will present many structurally and chemically equivalent motifs (*e*.*g*. many equivalent chain termini) for attachment or fusion – 12 for symmetry T, 24 for symmetry O, and 60 for symmetry I. That feature is beneficial for some applications (*e*.*g*. vaccine-like particles ^22–25^, polyvalent binding ^26,27^, enzymatic materials ^28,29^, and imaging scaffolds ^30–32^). However, for other applications it may be desirable to functionalize (or present fusions) on a singular location. It is notable that this type of ‘addressability’ is generally easy to achieve for nanomaterials based on nucleic acids, where symmetry is typically not a fundamental feature. One particular form of designed protein architectures – protein origami ^33^ – also lacks symmetry. Nonetheless, self-assembly design methods, whose successes are dramatically expanding due to modern machine learning algorithms ^7,13^, generally give rise to symmetric architectures, and thus to challenges in breaking symmetry for certain purposes.

One recent study succeeded in breaking the symmetry of a protein cage in order to produce a designed capsid larger than would otherwise be possible with purely icosahedral symmetry ^34^. That effort relied on an exceptionally complex and demanding task of designing a series of similar but selectively associating protein-protein interfaces. Fourteen sets of cage designs were required to identify successful cases of cages with broken symmetry. Motivated by the utility of protein cages with broken symmetry and by the difficulty associated with their creation, we sought a straightforward strategy that might allow the production of protein cages with broken symmetry and singular motifs for exterior binding or attachment.

In the present work, we demonstrate a new way to produce designed protein cages with broken symmetry. Our approach exploits a principle of steric collision in the protein cage interior to assure that only one copy of a modified subunit (bearing an extra inward-facing domain) can be accommodated in an otherwise symmetric cage. The asymmetry of the resulting cage is validated through biophysical and structural methods, including native mass spectrometry.

## RESULTS

### Protein design of a symmetry-broken cage

In designing a tetrahedral-type protein cage with broken symmetry, we took the previously characterized protein cage known as T33-51 as a starting point ^35^. T33-51 is a tetrahedrally symmetric architecture built from two subunit types, A and B, with stoichiometry A_12_B_12_. Four trimers of each subunit type sit on the body diagonals of a cube. The two naturally trimeric components of T33-51 spontaneously self-assemble into a supramolecular structure due to a binding association at a designed interface between them.

To develop a new approach for creating symmetry-broken cages, we introduced a modified protein component A, referred to as A’, which includes fusion to a small inward-facing protein domain. Owing to its inward-facing fusion, and the limited space in the interior of the protein cage, only one copy of the A’ subunit can be incorporated in an assembled cage without steric conflict; the other 11 copies must be occupied by the unmodified subunit A. A key motivation for breaking the symmetry of protein cages is to enable the display of unique attachments (not subject to symmetric repetition) on the exterior surface. Accordingly, in our design application, we further modified the A’ subunit with an outward-facing fusion to a DARPin protein domain. DARPin domains have been exploited as general binders, as their loop regions can be mutated (on the basis of library selection experiments) to bind diverse target proteins ^36–38^. Our fusion was to an anti-GFP DARPin, which has been the subject of recent protein cage applications ^30,31^. This design scheme is shown in Figure 1. To favor the desired assembly result, we co-transformed *Escherichia coli* with a gene for protein A’ in a low copy vector, with the A and B subunits of the T33-51 cage expressed on a high copy vector. We expected this scheme to generate a tetrahedral cage that displays a single anti-GFP DARPin on its surface (Figure 1a,b).

**Figure 1.**
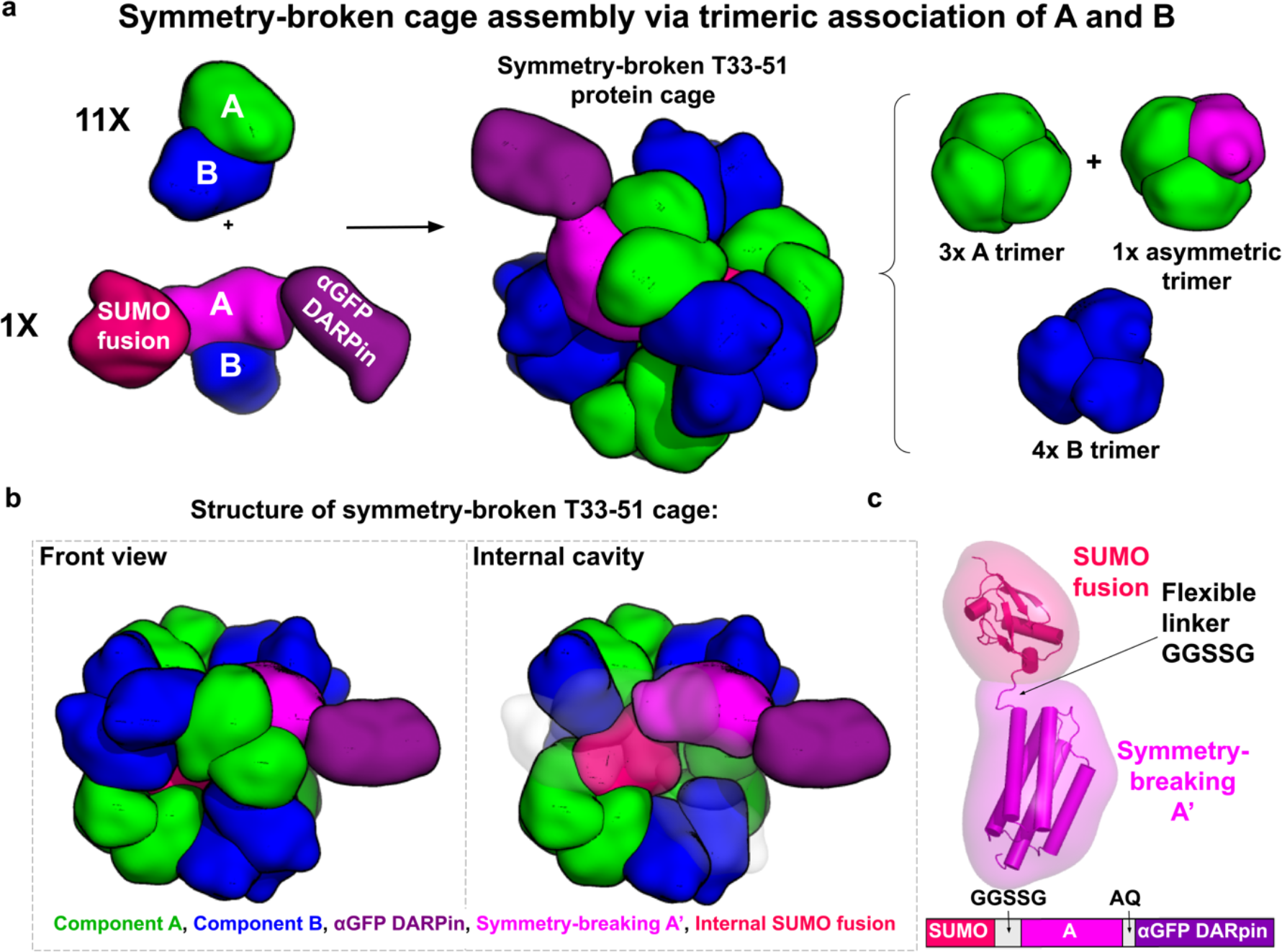
(a) Design of a symmetry-broken T33-51 protein cage. Subunits A (green) and B (blue) of the T33-51 cage were cloned into a high copy vector and co-transformed along with the A’ subunit (pink), which contains an internal SUMO fusion (red) and an outward-facing anti-GFP DARPin fusion (violet), in a low copy vector into *E. coli*. Due to its inward-facing SUMO fusion, and the limited space in the interior of the protein cage, only one copy of the A’ subunit can be incorporated in an assembled cage without steric conflict. **(b)** Structure of the symmetry-broken T33-51 cage. One A trimer contains a symmetry-broken A’ subunit (pink), which displays one anti-GFP DARPin (purple) on the cage surface and occupies the internal cavity of the cage with the inward-facing SUMO fusion (red). **(c)** Structure of symmetry breaking A’. In addition to an N-terminal (inward-facing) SUMO domain, the A’ subunit is fused to an outward-facing anti-GFP DARPin (not shown).

To evaluate inward-facing domains for fusing to subunit A’, we employed AlphaFold2 ^39^ as a convenient approach for obtaining plausible structural models. As candidates, we curated a set of common protein fusion tags, ranging from 10 to 20 kDa. We reasoned that this size range would allow for the fusion protein to fill a substantial fraction of the internal cavity of the cage without compromising its structural integrity. After reviewing these structure predictions, we selected the SUMO tag to serve as the internal fusion candidate based on its molecular weight, globular tertiary structure, and potential utility for subsequent characterizations. We evaluated models of the SUMO fusions with different linker lengths by aligning each version of the A’ subunit to the published crystal structure of the T33-51 cage ^35^ in PyMOL, noting whether the internal fusion protein fit within the cage interior without clashing with other A or B components. Based on this modeling, we chose GGSSG as a short flexible linker sequence (Figure 1c).

### Biochemical characterization of a symmetry-broken cage

We co-transformed the original A and B subunit genes for T33-51 in the high copy pRSFDuet-1 vector and the A’ gene in medium-to-low copy pET22b+ vector into *E. coli* BL21(DE3) cells. We purified the cages from cell lysate using immobilized nickel affinity chromatography based on a polyhistidine tail on the B subunit. SDS-PAGE analysis (Figure S1) confirmed the presence of the symmetry-broking A’ subunit in the presumptive protein cage complex obtained by nickel affinity purification.

Size exclusion chromatography (SEC) of the material obtained from nickel affinity purification confirmed successful cage assembly at the expected elution volume (Figure 2a), falling between that for the unmodified T33-51 cage and a T33-51 cage displaying 12 DARPins (one on each A component in the previous symmetric construction). When analyzed with blue-native PAGE, the SEC peak produced a distinct band at the expected molecular weight of a symmetry-broken cage (Figure 2b). SDS-PAGE analysis of the size exclusion chromatography fraction indicated the presence of symmetry-breaking component A’ in the cage peak fractions (Figure 2c), further confirming that the cage peak reflects symmetry-broken cages. Gel densitometry analysis of the bands corresponding to A, A’, and B subunit components confirmed the expected stoichiometric ratio between the A’ and A components (Figure 2d). In the selected SEC peak, the experimentally estimated ratio of A’ to A was 1.1:10.9, which is in close agreement with the theoretical value of 1:11, showing that only one copy of the A’ subunit was incorporated into the overall cage assembly, as expected.

**Figure 2.**
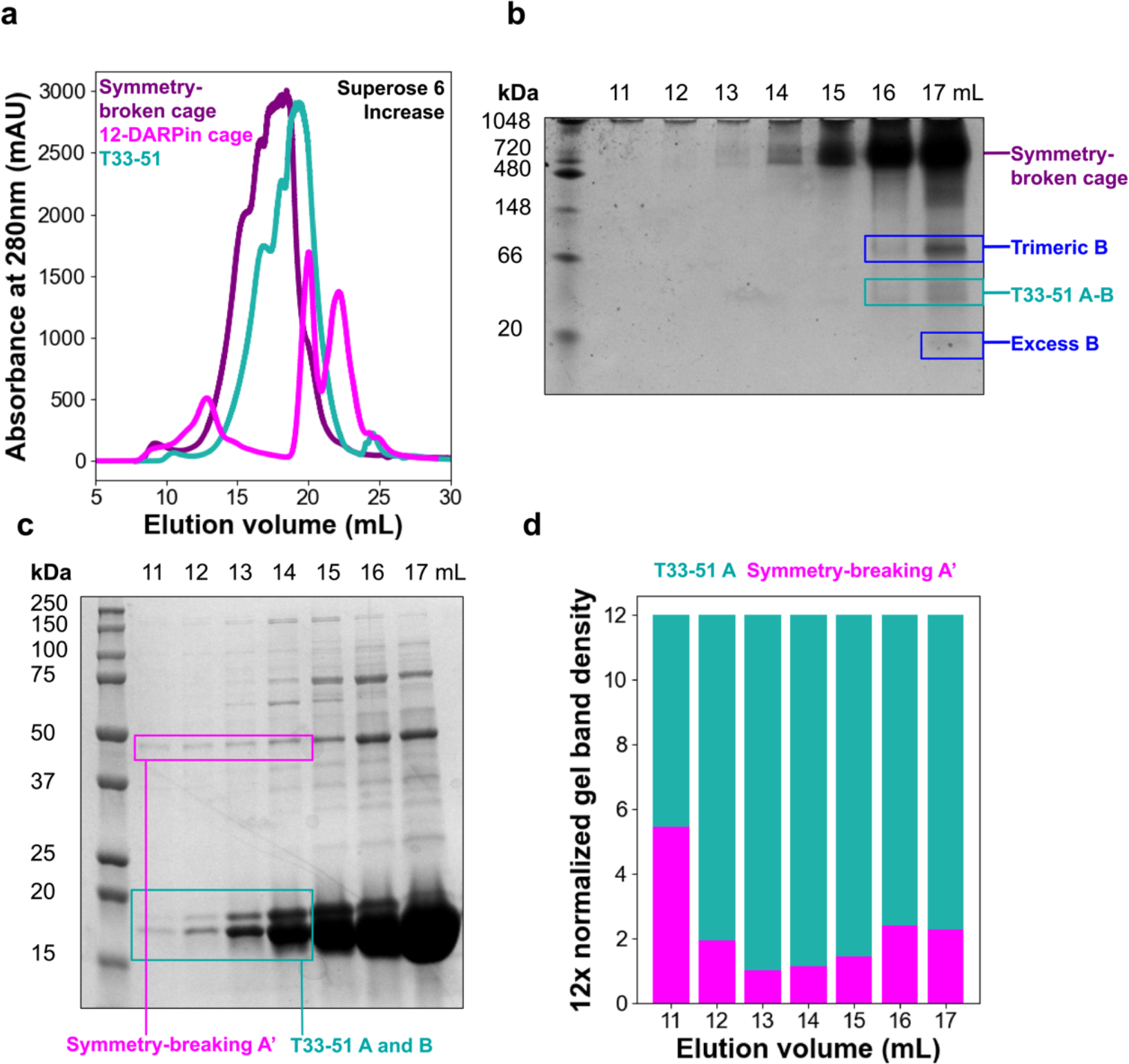
Purification of symmetry-broken T33-51 cage. **(a)** Size exclusion chromatography of immobilized nickel affinity chromatography-purified symmetry-broken cage (purple), unmodified T33-51 cage (teal) and T33-51 cage with surface anti-MBP DARPin fusions on all twelve A components (pink). Peaks corresponding to the cage appear from 12 to 14 mL. **(b)** Blue native PAGE of symmetry-broken cage size exclusion chromatography elution fractions (elution volume 11 mL to 17 mL). **(c)** SDS-PAGE of symmetry-broken cage size exclusion chromatography elution fractions (elution volume 11 mL to 17 mL). **(d)** Gel densitometry of each lane in (c). Molecular-weight normalized band density was multiplied by 12 to visualize more easily the stoichiometric ratio of T33-51 components to symmetry-broken A’ in each elution fraction.

### Structural characterization of the symmetry-broken cage

We pursued structural studies using electron microscopy and native mass spectroscopy. Through negative-strain microscopy (Figure 3a), we confirmed that the designed assembly forms geometrically regular particles of the expected size, reflective of intact cages. We also performed preliminary cryo-EM analysis. Here as well, images confirmed that intact cages were formed (Figure 3b). A low-resolution 3D reconstruction resulted in density closely matching the regular T33-51 cage. Not unexpectedly, image processing (even in the absence of symmetry constraints) did not reveal the singular protruding DARPin domain. Owing to its flexible attachment, small size, and 1/12 occupancy, particle alignment algorithms were not able to uniquely define the singular attachment. That finding provides an interesting counterpoint to recent cryo-EM scaffolding developments where symmetric and rigid attachments have proven critical for imaging ^30^.

**Figure 3.**
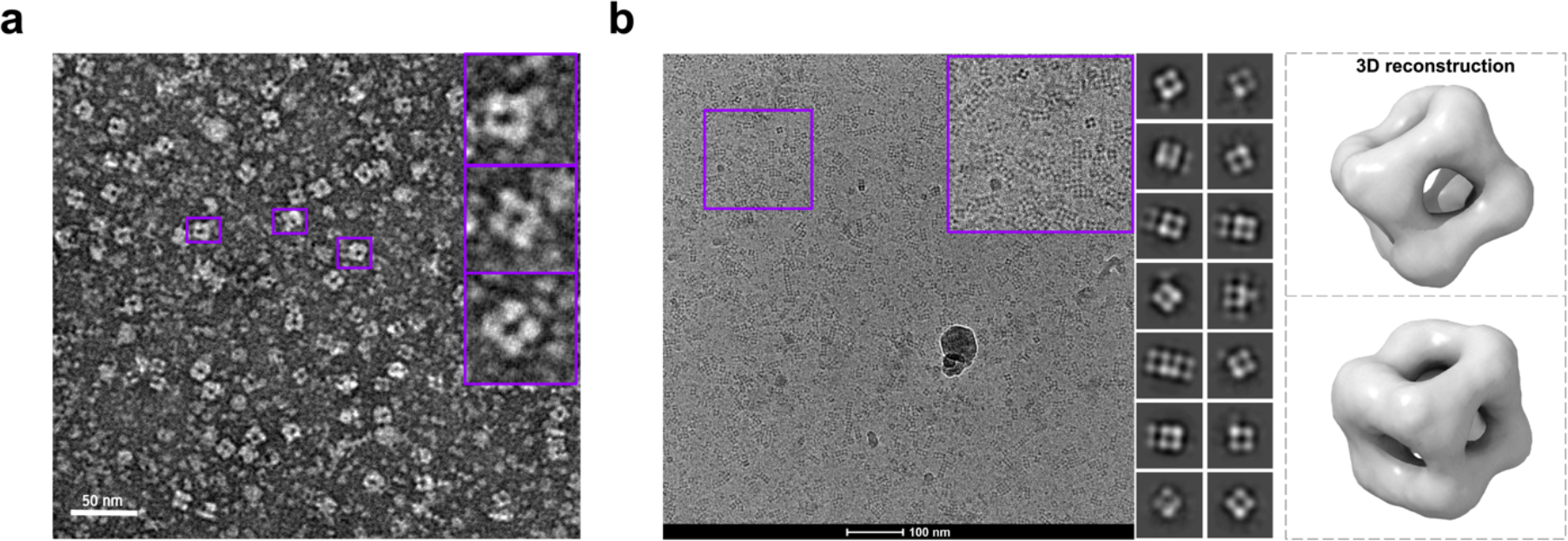
Structural characterization of the symmetry-broken cage. **(a)** Negative-stain electron microscopy of the symmetry-broken cage. Three particles of the expected size of a T33-51 cage that exhibit roughly cubic shape are highlighted. The scale bar is 50 nm. **(b)** Cryo-electron microscopy of the symmetry-broken cage. Twelve 2D particle classes are shown, which were used for the low-resolution 3D reconstruction (left) of the symmetry-broken cage core.

We turned to native mass spectrometry as a precise method for resolving the subunit stoichiometry of the assembled cages. Native mass spectrometry has proven useful in other studies on designed protein cages ^11,15,40–42^. Native mass spectrometry further confirmed the architecture and correct stoichiometry of the symmetry-broken cage (Figure 4). The dominant mass form (observed at ∼486 kDa) was within 0.2% of the calculated mass of a symmetry-broken cage with stoichiometry A11, A’1, B12 (485 kDa). Interestingly, the experimentally observed mass was slightly higher than the calculated model value. We hypothesize that this could reflect the incidental encapsulation of molecular species in the cell (where assembly occurs), and it is notable that a similar deviation was observed in prior work on hollow protein cages ^11^.

**Figure 4.**
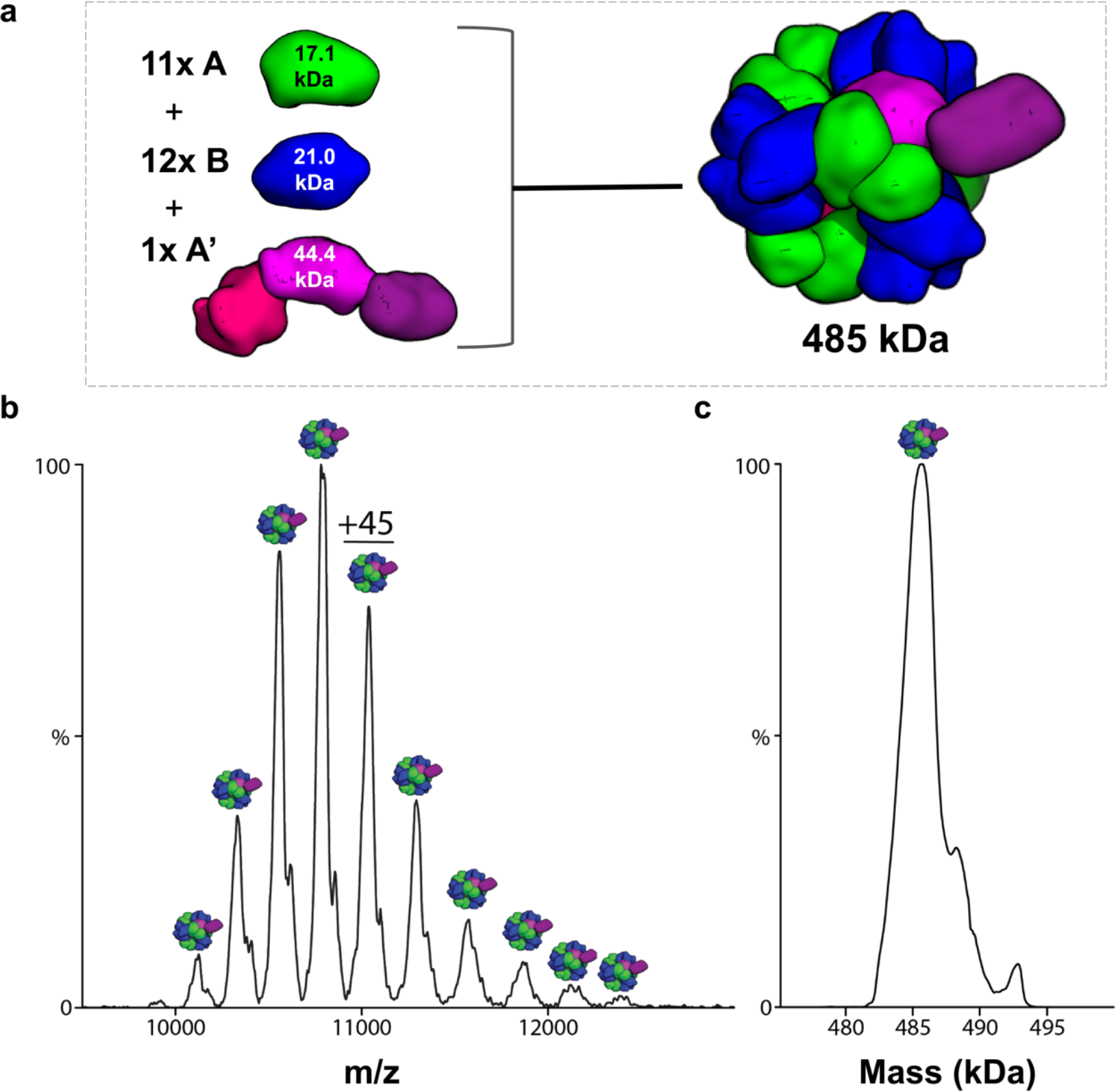
Native mass spectrometry of symmetry-broken cage. **(a)** The architecture of the symmetry-broken cage. The cage is composed of 11 copies of subunit A (17.1 kDa), 12 copies of subunit B (21.0 kDa), and one copy of subunit A’ (44.4 kDa). The total mass of the symmetry-broken cage is 485 kDa. **(b)** Native mass spectrum of symmetry-broken cage in 200 mM ammonium acetate. **(c)** Deconvolution of mass spectrum shown in (b).

## DISCUSSIONS AND CONCLUSION

The central aim of this work was to demonstrate a method for generating symmetry-broken protein cages. Our approach, exploiting a strategy of steric occlusion, was successful in that goal. A modified version of an otherwise tetrahedrally symmetric cage was shown to comprise a single copy of a unique subunit variant, thereby allowing the presentation of a single copy of an outward-facing binding domain. The relative simplicity of the present design approach is notable. A suitable protein cage having a subunit with an inward-facing terminus is required, and an appropriate choice must be made for the interior occluding domain, but no other challenging design elements (e.g., novel protein-protein interfaces) were required to achieve the desired goal.

As designed protein cages find increasing uses in nanotechnology and medicine, the ability to prepare asymmetric or addressable variants will become more important. Symmetric versions of protein cages have proven critically useful in single-particle cryo-EM applications as imaging scaffolds ^30–32^, but asymmetric forms could be advantageous in other imaging applications, including in *in-situ* cryo-EM tomography, where it might be desirable to avoid the effects of high-copy number binding in a native cellular environment. Accordingly, symmetry-broken cages with singular binding motifs on their exterior could prove valuable as mid-nanometer, targeted geometric markers inside cells. Those applications are currently under investigation.

## METHODS

### Design of symmetry-breaking component A’

A set of common fusion tags was curated and evaluated for suitable size (<20 kDa) and possible utility in the context of cage characterization to serve as internal fusions for the symmetry-breaking component A’. From this set, the small ubiquitin-like modifier (SUMO) protein was selected. Next, a set of linker sequences, which included the short linker AQ, the flexible linker GGSSG, the rigid linker EEEAQKAA, and several alpha-helical linkers between A and the internal fusion were evaluated for the SUMO tag in order to determine which linker would afford the most favorable placement of the internal fusion relative to A and the cage cavity. This evaluation was performed by obtaining structure predictions of the A-linker-internal fusion protein using AlphaFold2 ^39^ and aligning this prediction to the published crystal structure of the T33-51 cage (PDB ID: 5cy5) in PyMOL ^35^. The final chosen design for the symmetry-breaking component A’ used the linker GGSSG.

### Cloning and protein expression

A DNA fragment containing the sequence for T33-51 components A and B separated by a spacer (sequence shown below) sourced from Cannon *et al*. (2020) was synthesized (Twist Biosciences) and cloned into the pRSFDuet-1 vector (gifted by Dr. Mark Arbing, UCLA-DOE IGP) using Gibson assembly cloning. Separately, DNA fragments containing the four chosen symmetry-breaking A designs were synthesized (Twist Biosciences) and cloned into the pET22b+ vector via Gibson assembly. Plasmid DNA for all constructs was cloned in *E. coli* DH5alpha cells (New England Biolabs) and extracted using ZymoPURE Plasmid Miniprep Kits.

Purified T33-51 and symmetry-breaking A constructs were co-transformed into *E. coli* BL21(DE3) cells and plated on Luria Bertani broth (LB) agar supplemented with 50 μg/mL kanamycin and 50 μg/mL ampicillin. Three colonies from each plate were used to inoculate 50 mL of LB supplemented with 50 μg/mL kanamycin and 50 μg/mL ampicillin. The LB growths were then incubated with shaking overnight at 37°C and 200 rpm. Following overnight incubation, 10 mL of each growth was used to inoculate 1L of kanamycin and ampicillin-supplemented LB. Growths were incubated with shaking at 37°C and 180 rpm until they reached an OD_600_ of 0.6, at which point expression was induced with 100 μM isopropyl β-d-1-thiogalactopyranoside. Growths were incubated with shaking at 18°C and 180 rpm for 18 hours, after which cells were harvested by centrifugation at 4000 xg for 20 minutes at 4°C.

### Protein purification

Harvested cell pellets were resuspended in lysis buffer (250 mM NaCl, 50 mM Tris-HCl pH 8.0, 20 mM imidazole pH 8.0, 2% w/v glycerol) supplemented with lysozyme, EDTA-free protease inhibitor cocktail, DNAse, and RNAse (Thermo Fisher Scientific). Cells were then lysed using an Avestin EmulsiFlex C3 homogenizer. Cell lysates were clarified for 30 minutes at 20,000 xg at 4°C. The clarified cell lysates were loaded onto a column containing equilibrated immobilized nickel affinity chromatography resin (Thermo Fisher Scientific) and eluted using a linear imidazole gradient from 50 mM to 250 mM. All immobilized nickel chromatography fractions, along with the cell pellet, were analyzed using SDS-PAGE.

100 mM imidazole and 250 mM imidazole elution fractions were concentrated using Millipore Amicon Ultra 100 kDa molecular weight cutoff filters at 3500 xg and 4°C. Concentrated protein samples were further purified via size-exclusion chromatography using a Superose 6 Increase column (GE Biosciences) and eluted with SEC buffer (250 mM NaCl, 50 mM Tris-HCl pH 8.0, 2% w/v glycerol). Size exclusion chromatography fractions corresponding to suspected cage peaks were analyzed via SDS-PAGE and native (non-denaturing) PAGE.

### Negative stain electron microscopy

Following size exclusion chromatography, symmetry-broken cage samples were diluted to 50 μg/μL. Formvar/Carbon 400 mesh copper grids (Ted Pella Inc) were glow-discharged for 30 seconds at 15 mA using a PELCO EasiGlow. 5 μL of cage sample was transferred to the glow-discharged grids. After 30 seconds of incubation, the sample was removed by blotting, and the grid was stained with 2% uranyl acetate for 30 seconds. The grid was then imaged using a Tecnai T12 electron microscope.

### Cryo-electron microscopy

Following size exclusion chromatography, symmetry-broken cage samples were concentrated and mixed with purified GFP in a final ratio of 1:12 cage to GFP at 0.6 mg/mL final cage concentration. 3.5 μL of the sample mixture was applied to glow discharged Quantifoil 300 mesh R2/2 copper grids and frozen with liquid ethane using a Vitrobot Mark IV (FEI). Grids were imaged using a Talos F200C. Automatic particle picking, 2D classification, and 3D reconstruction were performed using CryoSPARC ^43^.

### Native PAGE

Following size exclusion chromatography, fractions spanning the obtained cage peak were sampled and diluted 4:1 with ThermoFisher NativePAGE Sample Buffer (4X). The native PAGE was run at 150V for 105 minutes at 4°C in ThermoFisher NativePAGE Running Buffer.

### Native Mass Spectrometry

Size exclusion chromatography fractions corresponding to the symmetry-broken cage peak were pooled and concentrated to 50 μM. Samples were buffer-exchanged using Micro Bio-Spin 6 desalting column (Bio-Rad) into 200mM ammonium acetate (pH adjusted to 7.4 with ammonium hydroxide). Samples were diluted to 2 μM and loaded into pulled borosilicate glass capillaries prepared in-house. Samples were electrosprayed with the voltage applied using a platinum wire directly inserted into the solution into a Q Exactive Ultra-High Mass Range Hybrid Quadrupole-Orbitrap mass spectrometer (Thermo Scientific). Instrument resolution was set to 3125, in-source collision energy dissociation at 30V, collision energy at 50V, source temperature at 170°C, capillary voltage at 1.3kV, source DC offset at 21V, inter flatapole lens at 10V, injection flatapole DC at 15V, bent flatapole DC at 15V, transfer multiple DC at 1V, and trapping gas pressure at 7.5.

### Figure visualization

Symmetry-broken cage structures were created using published crystal structures of T33-51 A and B components (PDB ID: 5cy5) ^35^ and AlphaFold2 predictions for the symmetry-breaking A with the SUMO tag and short linker. These structures were rendered in PyMOL (Version 2.0 Schrödinger, LLC). Gel densitometry analyses were carried out using the Gel Analyzer feature in ImageJ ^44^.

## Acknowledgments

The authors thank Mark Arbing for providing the pRSFDuet-1 vector and for helpful discussions regarding cloning. We also thank Duilio Cascio and Alex Lisker for computing support.

