## Supplementary Material for "Design of a symmetry-broken tetrahedral protein cage by a method of internal steric occlusion"

Gladkov, *et al.*

**Supplementary Figure:**

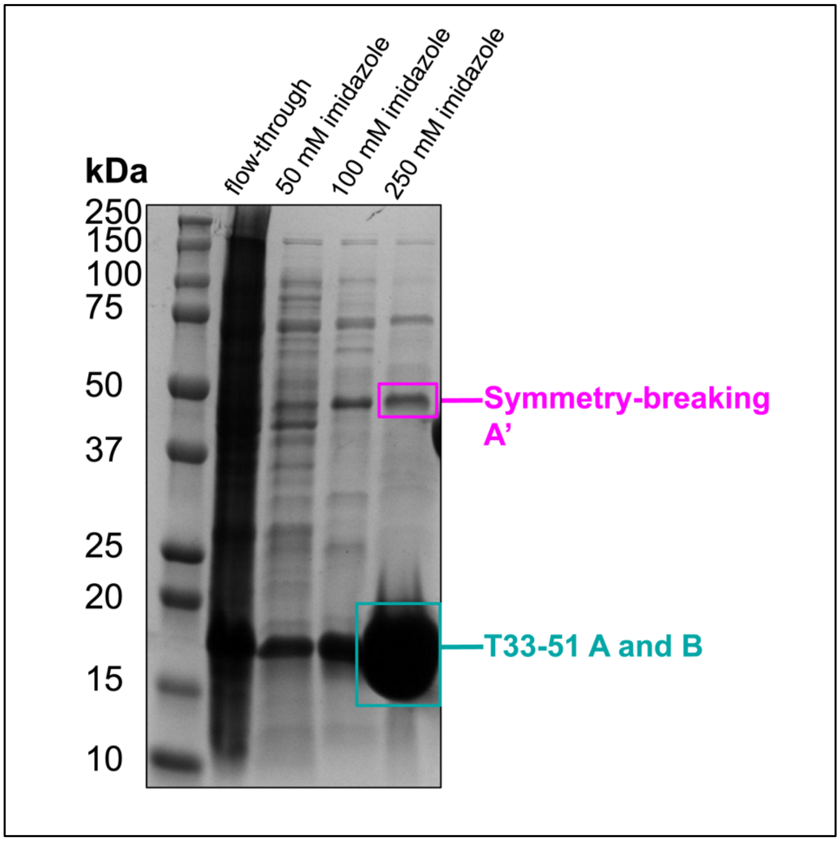


**Figure S1.** SDS-PAGE of immobilized nickel affinity chromatography fractions from the purification of symmetry-broken cage eluted with a linear imidazole gradient from 50 mM to 250 mM.

**Supplementary Text:**

Protein sequences:

| | |
|---|---|
| T33-51 A | MSNEEVWKDDPIIEANGTLDELTSFIGEAKHYVDEEMKGILEEIQNDIY<br>KIMGEIGSKGKIEGISEERIKWLAGLIERYSEMVNKLSFVLPGGTLESA<br>KLDVCRTIARRAERKVATVLREFGIGTLAAIYLALLSRLLFLLARVIEIEK<br>NKL |
| T33-51 A’ | MSDSEVNQEAKPEVKPEVKPETHINLKVSDGSSEIFFKIKKTTPLRRL<br>MEAFAKRQGKEMDSLRFLYDGIRIQADQTPEDLDMEGGSSGSNEEV<br>WKDDPIIEANGTLDELTSFIGEAKHYVDEEMKGILEEIQNDIYKIMGEI<br>GSKGKIEGISEERIKWLAGLIERYSEMVNKLSFVLPGGTLESAKLDVC<br>RTIARRAERKVATVLREFGIGTLAAIYLALLSRLLFLLARVIEIEKNKLGP<br>GSDLGKKLLEAARAGQDDEVRILMANGADVNAADDVGVTPLHLAAQ<br>RGHLEIVEVLLKYGADVNAADLWGQTPLHLAATAGHLEIVEVLLKNG<br>ADVNARDNIGHTPLHLAAWAGHLEIVEVLLKYGADVNAQDKFGKTPF<br>DLAIDNGNEDIAEVLQKAA |
| T33-51 B | MFTRRGDQGETDLANRARVGKDSPVVEVQGTIDELNSFIGYALVLSR<br>WDDIRNDLFRIQNDLFVLGEDVSTGGKGRTVTMDMIIYLIKRAVEMKA<br>EIGKIELFVVPGGSVESASLHMARAVSRRLERRIKAASELTEINANVLL<br>YANMLSNILFMHALISNKRLNIPEKIWSIHRVSLEHHHHHH |
